# Imaging photosynthetic pH dynamics in living plants and algae

**DOI:** 10.64898/2026.11.23.768545

**Authors:** Ana Paula Cislaghi, Minh Hoang, Pedro Barreto, Anna Erpenbeck, Ke Zheng, Laura Fischer, Eduard Windenbach, Sarah Käbe, Marlene Elsässer, Ute Armbruster, Hans-Henning Kunz, Michael Schroda, Michael Hippler, Felix Buchert, Karin B. Busch, Markus Schwarzländer

## Abstract

Photosynthetic light reactions generate a proton motive force across the thylakoid membrane, composed of a pH gradient (ΔpH) and an electric potential (Δψ). However, non-invasive dynamic tracking of the pH gradient across chloroplast sub-compartments has remained technically challenging *in vivo*. Here, we systematically evaluated four genetically encoded pH biosensors, cpYFP, pHGFP, Lime and pHluorin2, *in vitro* and *in planta* to provide a versatile pH-sensing toolbox. pHluorin2 combined a ratiometric readout, high brightness and responsiveness in the acidic range, and was selected for further work. In *Arabidopsis thaliana*, stromal pHluorin2 resolved rapid light-driven alkalinization with a light intensity-dependent transient overshoot, mirrored by a transient acidification reported by lumen-targeted pHluorin2. In *Chlamydomonas reinhardtii*, stromal pH rose gradually without overshoot and declined slowly in the dark, pointing to distinct regulation of stromal pH in algae and land plants. Pharmacological and genetic perturbation of electron transport and ΔpH indicated that pHluorin2 specifically reports photosynthesis-driven pH changes. As such our toolbox enables real-time, compartment-specific, high-resolution quantitative imaging to study photosynthetic proton dynamics in land plants and algae.

## Introduction

Photosynthesis converts the physical energy of sunlight into the chemical energy that sustains nearly all forms of life. This conversion relies on the transmembrane proton motive force (*pmf*) as a universal intermediate, which is generated by light-driven electron transport across the photosynthetic membrane (Kramer et al., 2003). The *pmf* has two components: a transmembrane pH gradient (ΔpH) and an electrical potential (Δψ). Either component can drive ATP synthesis via thylakoid ATP synthase (Junge & Nelson, 2015; Kramer et al., 2003), but they fulfill distinct regulatory roles in the chloroplasts of plants and algae. ΔpH is generally considered the dominant component in magnitude (Wilson et al., 2021) and has major regulatory importance.

In land plants, lumen acidification protonates photosystem II subunit S (PsbS) and activates violaxanthin de-epoxidase, triggering qE, the fastest component of non-photochemical quenching (NPQ) (Avenson et al., 2005; Li et al., 2000; Ruban, 2016). It also slows plastoquinol oxidation at the cytochrome _b6f_ complex (Rumberg & Siggel, 1969; Munekage et al., 2001), which protects photosystem I from over-reduction under fluctuating light (Joliot & Johnson, 2011; Suorsa et al., 2012), and the accompanying alkalinization of the stroma regulates Calvin–Benson cycle enzymes (Werdan et al., 1975). In contrast, Δψ does not trigger these regulatory responses, and an excessive Δψ can even promote charge recombination at photosystem II, causing photodamage (Davis et al., 2016). In the green alga *Chlamydomonas reinhardtii*, ΔpH also controls photoprotection and electron transport, but through partly different components. Lumen acidification triggers qE by protonation of the light-harvesting complex stress-related protein 3 (LHCSR3) instead of PsbS (Peers et al., 2009; Ballottari et al., 2016), and electron transfer at the cytochrome b_6_f complex is slowed under acidic lumenal conditions (Ozawa et al., 2023). Chloroplasts host regulatory systems to actively adjust ΔpH and match metabolic demands by modulating ATP synthase conductivity (Kanazawa & Kramer, 2002; Rott et al., 2011) as well as through regulating ion fluxes. Key ion fluxes across the thylakoid membrane include those mediated by the K^+^/H^+^ antiporter K^+^ EFFLUX ANTIPORTER 3 (KEA3) and the voltage-dependent chloride channel 1 (VCCN1), both of which shape *pmf* partitioning between ΔpH and Δψ (Armbruster et al., 2014; Herdean et al., 2016; Kunz et al., 2014). Measuring pH dynamics in the stroma and the thylakoid lumen *in vivo* with high spatial and temporal resolution therefore holds the potential to resolve the kinetics of ΔpH build-up and dissipation at dark–light transitions in specific tissues, cells, individual chloroplasts and even thylakoid sub-systems, how electron flow and ion fluxes shape stromal and luminal pH, and how these dynamics differ between plants and algae.

Several techniques have been used to estimate luminal and stromal pH. The electrochromic shift (ECS) assay reports changes in the transthylakoid electric field, and its signal kinetics are used to derive ΔpH indirectly, using assumptions that have been controversially debated (Cruz et al., 2001; Kramer et al., 2003; Johnson & Ruban, 2014; Bailleul et al., 2010; Allorent et al., 2018). To determine lumen acidification, quenching of chlorophyll fluorescence is widely used as a proxy (Briantais et al., 1979; Horton et al., 1991). Yet, quenching reports the response of the antenna, which also depends on parameters other than pH alone, such as PsbS or LHCSR3 content, xanthophyll de-epoxidation and physiological conditions (Kanazawa & Kramer, 2002; Li et al., 2002; Peers et al., 2009). Neither approach measures pH directly, and direct methods have remained largely restricted to isolated chloroplasts or thylakoids. These include accumulation of the fluorescent dye 9-aminoacridine, which, however, has limited sensitivity at low ΔpH and is prone to artefacts due to membrane binding (Schuldiner et al., 1972; Su & Lai, 2017). Moreover, the distribution of radio-labelled weak acids and bases can be exploited, for pH assessment of the thylakoid lumen and stroma. However, this method requires cumbersome sampling for fractionation which limits the temporal resolution that can realistically be achieved (Heldt et al., 1973). Other approaches use pH-sensitive nitroxide spin probes detected by electron paramagnetic resonance (EPR) for the lumen (Tikhonov et al., 2008) and the fluorescent dye BCECF-AM for the stromal pH measurements (Rink et al., 1982; Su & Lai, 2017). Overall, and despite their individual limits, these approaches have been instrumental for our current understanding of *pmf*-related pH dynamics. None of them has enabled dynamic *in vivo* pH measurements in both the stroma and the thylakoid lumen of chloroplasts in intact living cells, however. As a result, there remains a major void in our understanding of photosynthesis-associated pH as one of the central energetic hubs in biology.

Genetically encoded pH biosensors are fluorescent proteins whose spectral properties change depending on pH in their immediate environment (Bizzarri et al., 2009; Miesenböck et al., 1998). These pH sensors have the potential to address several of the remaining limitations. They can be targeted genetically to specific cell (sub-)compartments and used non-invasively in intact tissue based on their fluorescence properties. Several pH biosensors show ratiometric spectral changes that make the detected signal largely independent of sensor concentration, i.e. expression levels of the fluorescent protein. Further, pH values can be estimated from the ratiometric fluorescence values through *in situ* calibration. Such pH biosensors were shown to resolve light-induced stromal alkalinization in *Arabidopsis* leaves (Elsässer et al., 2020; Zheng et al., 2025; Loro et al., 2016; Uflewski et al., 2024), and luminal acidification in isolated chloroplasts (Yang et al., 2017). However, *in vivo* pH measurement in the thylakoid lumen has not been replicated since and has remained limited by reliable sensor expression and targeting, as well as the need for sensors that provide an informative read-out at low pH. Biosensors differ in their pKa, and hence the pH range over which they respond, and in their spectroscopic dynamic range, which together contribute to distinct sensitivities of pH biosensor variants. Further, their colour and brightness (i.e. quantum yield and levels of *in vivo* expression) are critical considerations for measurements in the presence of chlorophyll. Hence, the careful selection and validation of suitable sensors remains a critical step towards resolving photosynthesis-driven pH dynamics *in vivo*.

Here, we introduce a toolkit of four pH biosensor variants for measuring *in vivo* pH dynamics in the chloroplast stroma of the plant model *Arabidopsis thaliana* and the algae model *Chlamydomonas reinhardtii.* We empirically compare and validate the properties of the different sensor systems in their response to illumination-induced pH changes, based on which we make specific and practical recommendations for their deployment in photosynthesis research. We add a luminal pH sensor as a novel resource, to pave the way towards ΔpH measurements *in vivo*, scalable from chloroplast subdomains up to whole cells, tissue-types, organs (most importantly leaves) and plants.

## Results

### Spectroscopic profiling and titration of purified GFP-derived pH biosensors

We selected established fluorescent protein-based pH biosensors with emission distinct from chlorophyll fluorescence. Furthermore, to maximize the odds of efficient import across the chloroplast membrane systems, we restricted the selection to sensors consisting of a single fluorescent protein (Niemeier et al., 2026). Based on those considerations we focused on four pH sensors for systematic side-by-side assessment: cpYFP (Baird et al., 1999; Schwarzländer et al., 2014), pHGFP (Moseyko & Feldman, 2001), Lime (Shen et al., 2023) and pHluorin2 (Mahon, 2011). cpYFP, pHGFP and pHluorin2 are dual-excitation ratiometric sensors, whereas Lime responds mainly at one excitation wavelength (Fig. 1b). All four derive from wild-type *Aequorea victoria* GFP (avGFP) through distinct engineering routes, including circular permutation, chromophore-proximal substitutions and stabilizing mutations (Fig. 1a). We compared the excitation and emission spectra of the purified sensor proteins *in vitro* across a broad, physiologically relevant pH range (Fig. 1b and Fig. S1).

**Figure 1.**
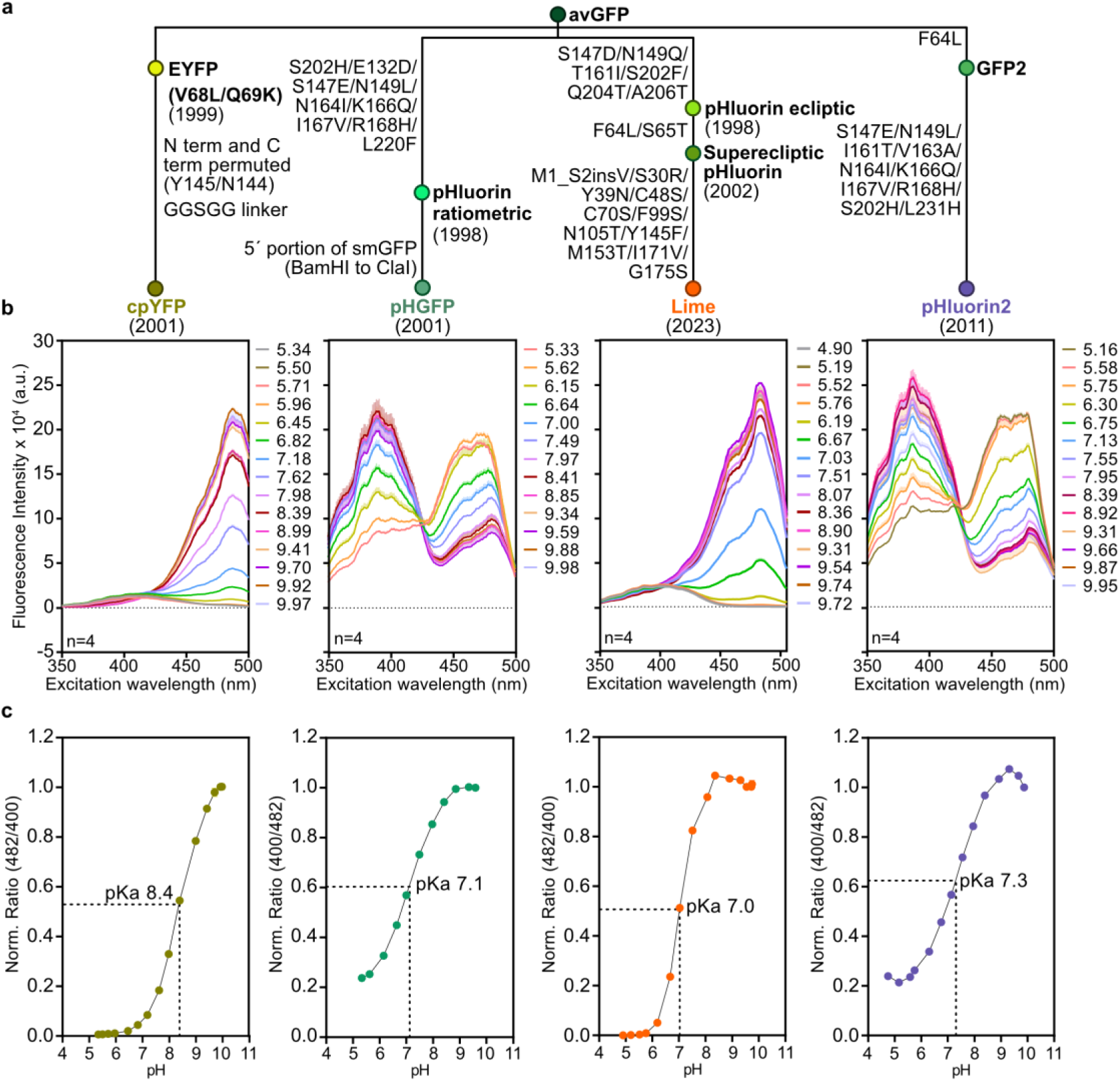
*In vitro* characterization of four GFP-derived pH sensors. **a.** Derivation of cpYFP, pHGFP, Lime and pHluorin2 from wild-type *Aequorea victoria* GFP (avGFP), with the engineering steps indicated along the branches and the year of the original publication of each sensor. **b.** Excitation spectra (350–500 nm; emission at 510 nm) of purified recombinant cpYFP, pHGFP, Lime and pHluorin2 in universal buffer at pH ∼5–10. Lines, mean of n = 4 technical replicates. **c.** Ratio of the excitation intensities at 482 and 400 nm (cpYFP, Lime) or 400 and 482 nm (pHGFP, pHluorin2), normalized to the maximum, as a function of pH. Solid lines, sigmoidal fits used to obtain the apparent pKa; dashed lines, apparent pKa (cpYFP 8.4, pHGFP 7.1, Lime 7.0, pHluorin2 7.3); symbols, mean ± SD of n = 4 technical replicates (four wells of the same protein preparation).

Excitation spectra recorded between pH ∼5 and 10 showed pH-dependent responses (Fig. 1b) consistent with previous reports for each of the sensors (Miesenböck et al., 1998; Schwarzländer et al., 2011; Mahon, 2011; Shen et al., 2023). With increasing pH, fluorescence excited at ∼482 nm increased for cpYFP and Lime, whereas pHGFP and pHluorin2 showed a reciprocal switch, with fluorescence excited at ∼390 nm increasing and at ∼482 nm decreasing. Emission spectra were similar for all four sensors, with a single maximum at ∼510 nm upon excitation at either 405 or 482 nm (Fig. S1). Two properties are most relevant for measuring pH in chloroplasts: emission at ∼510 nm, which is spectrally well separated from chlorophyll fluorescence, and excitation at ∼400 and ∼482 nm, which allows one to determine the emission intensity ratio after excitation with the two distinct excitation wavelengths (482/400 for cpYFP and Lime; 400/482 for pHGFP and pHluorin2) as a readout for pH, largely independently of sensor concentration. We therefore used this ratio to determine the pKa, the pH response range and the fold change of each purified biosensor in our empirical side-by-side comparison.

The pH titrations showed a sigmoidal relationship between spectroscopic ratio and pH for all sensors (Fig. 1c; Table S1). cpYFP had the highest apparent pKa (8.4), consistent with the pKa of ∼8.7 estimated for the purified sensor (Schwarzländer et al., 2014), followed by pHluorin2 (7.3), pHGFP (7.1) and Lime (7.0). The pKa values of Lime and pHGFP were close to those reported for the parental sensors (7.1 for Lime, Shen et al., 2023; 6.9 for ratiometric pHluorin, from which pHGFP is derived, Miesenböck et al., 1998), whereas that of pHluorin2 was ∼0.4 units above the previously reported ∼6.9. Lime had a steeper transition than the other sensors (Hill coefficient, nH, of 1.38 versus 0.74–0.85), resulting in a narrower window with a close-to-linear response. The pH response ranges, calculated as the pH interval between 10% and 90% of the maximal ratio response, were 7.3-9.5 for cpYFP, 5.8-8.4 for pHGFP, 6.4-7.7 for Lime and 5.9-8.5 for pHluorin2 (Table S1). Because the stromal pH is expected to show shifts within the range of pH 7 to 8 during illumination (Heldt et al., 1973; Su & Lai, 2017; Shen et al., 2013), we compared the sensors in this range: between pH 7 and 8, the ratio changed 5.1-fold for cpYFP, 2.0-fold for Lime, 1.7-fold for pHluorin2 and 1.5-fold for pHGFP (Table S1). At pH 8.0, cpYFP had reached only 32% of its maximal response, whereas Lime (96%), pHGFP (82%) and pHluorin2 (78%) were closer to saturation. The acidic range between pH 5.5 and 7 as relevant for the lumen (Heldt et al., 1973; Takizawa et al., 2007), showed the opposite: the ratio of pHGFP and pHluorin2 changed 2.3- and 2.1-fold, respectively, whereas cpYFP and Lime responded too little at pH 5.5 for the fold change to be determined. Among the four empirically and side-by-side assessed pH biosensors, pHGFP and pHluorin2 therefore are the most appropriate candidates for luminal pH sensing, whereas, in the *in vitro* titrations, cpYFP appeared best suited to resolve pH dynamics in the alkaline range of the stroma.

### Subcellular targeting and spectral properties in Arabidopsis leaves

To compare the sensors *in planta*, we generated constructs for stromal expression of pHGFP, pHluorin2 and Lime. Each sensor was fused to the chloroplast transit peptide of NADPH-dependent thioredoxin reductase C (NTRC; Serrato et al., 2004). pHGFP was expressed under the control of the *UBIQUITIN 10* promoter (*UBI10*; Norris et al., 1993; Grefen et al., 2010), and pHluorin2 and Lime under the control of the *ASPARTIC PROTEASE A1* promoter (*APA*; D’Ippólito et al., 2020) (Fig. 2a). For stromal cpYFP, we used lines generated previously (Behera et al., 2018), in which cpYFP is fused to the transketolase transit peptide (TKTP) and expressed under the *Cauliflower mosaic virus 35S* promoter (*CaMV35S*). Following *Agrobacterium*-mediated transformation of Arabidopsis and selection of stable transformants, we verified sensor expression in leaf tissue using two-dimensional excitation–emission fluorescence spectroscopy (2D-EEM; Fig. 2b). Non-transformed Col-0 leaves displayed moderate intrinsic fluorescence at excitation ∼350 nm and emission 425–475 nm and from chlorophyll at emission 650–700 nm across a broad excitation range (Donaldson, 2020). Importantly, the signals of all four sensors were distinct from these areas of intrinsic fluorescence. Yet, fluorescence intensity from the biosensors differed markedly among the transformants: cpYFP lines exhibited a comparatively weak signal at 488 nm excitation; pHGFP lines showed a moderate signal across ∼400 to ∼488 nm excitation; Lime lines showed a strong signal centered around ∼488 nm excitation; and pHluorin2 lines showed the highest overall signal, with a broad excitation range around ∼400 and ∼488 nm similar to that of pHGFP (Fig. 2b). While these differences are affected by sensor quantum yield, they are largely a result of expression and accumulation of the mature sensor protein *in planta*.

**Figure 2.**
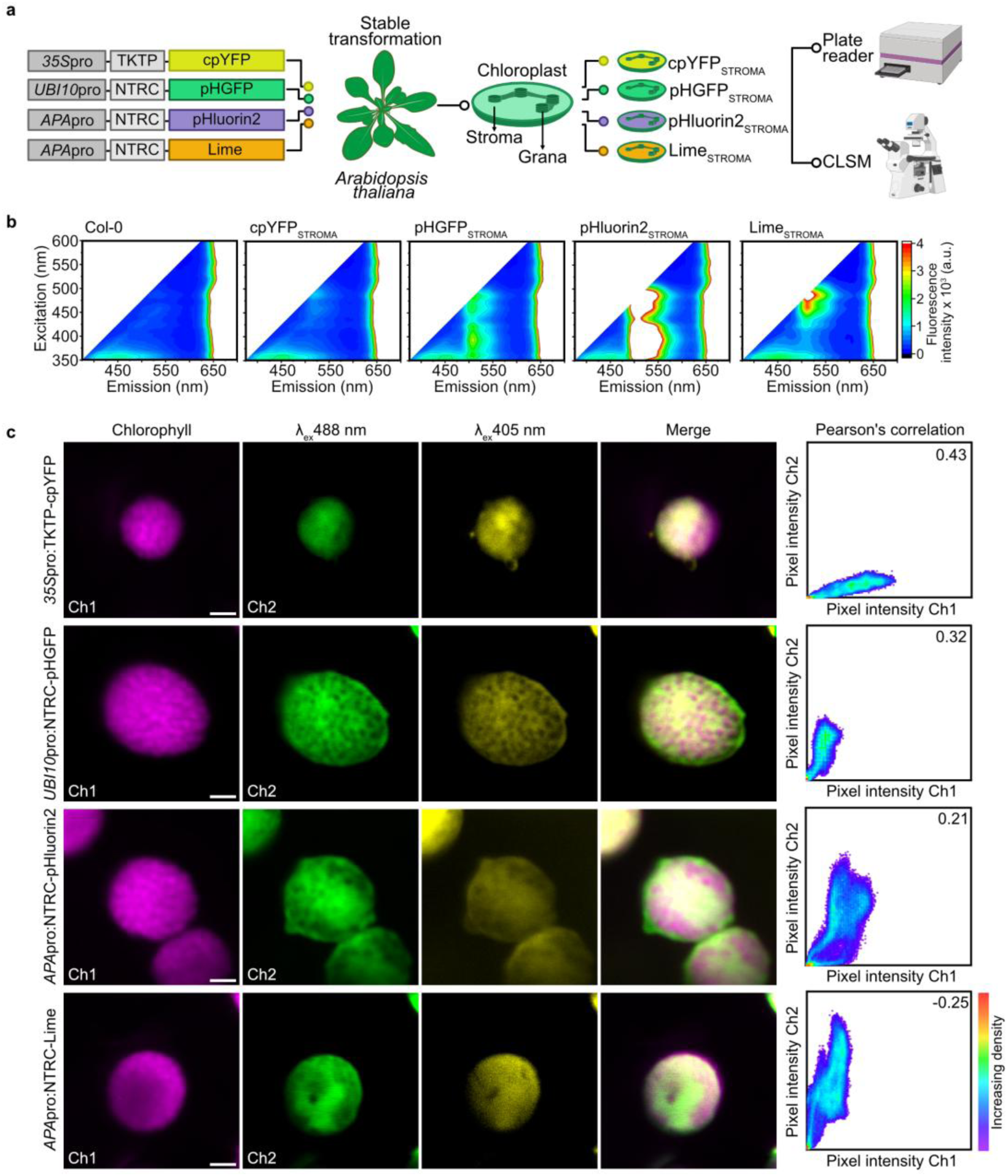
Chloroplast stroma-targeted pH sensors in Arabidopsis. **a.** Constructs used to generate stable *Arabidopsis thaliana* lines expressing cpYFP, pHGFP, pHluorin2 or Lime in the chloroplast stroma: *35S*pro:TKTP-cpYFP, *UBI10*pro:NTRC-pHGFP, *APA*pro:NTRC-pHluorin2 and *APA*pro:NTRC-Lime. *35S*, *Cauliflower mosaic virus 35S* promoter; *UBI10*, *UBIQUITIN 10* promoter; *APA*, *ASPARTIC PROTEASE A1* promoter; TKTP, transketolase transit peptide; NTRC, NADPH-dependent thioredoxin reductase C transit peptide. The plate reader and confocal microscope icons were created with BioRender.com (Schwarzlander, M. (2026) https://BioRender.com/8ypyqho). **b.** Two-dimensional excitation–emission matrices (2D-EEM) of leaf discs from Col-0 and from each sensor line, recorded on a plate reader (excitation 350–600 nm; emission up to 700 nm) with identical gain settings. The colour scale shows fluorescence intensity (arbitrary units × 10^3^): from 0 (black) to 4 (red); white areas, signal above the maximum of the scale. A representative scan of three leaf discs per genotype is shown. **c.** Confocal laser scanning microscope (CLSM) images of mesophyll chloroplasts showing chlorophyll fluorescence (Ch1, magenta; excitation 405 nm, emission 655–735 nm), sensor fluorescence excited at 488 nm (Ch2, green; emission 495–550 nm) and at 405 nm (yellow; emission 495–550 nm), and the merge. Right, pixel-intensity scatter plots of Ch2 versus Ch1 (colour, pixel density) with Pearson’s correlation coefficients (cpYFP 0.43, pHGFP 0.32, pHluorin2 0.21, Lime −0.25), calculated within a region of interest covering the whole chloroplast. Scale bars, 2 µm.

We next examined the sub-cellular and sub-organellar localization of the sensors in leaf mesophyll (Fig. 2c) and epidermal cells (Fig. S2) using confocal laser scanning microscopy (CLSM). In all four lines, sensor fluorescence exclusively filled the chloroplast interior and was distinct from the discrete, high-intensity chlorophyll fluorescence in grana thylakoid membranes. Pearson’s correlation coefficients between sensor and chlorophyll fluorescence were low or negative (cpYFP: 0.43; pHGFP: 0.32; pHluorin2: 0.21; Lime: −0.25; Fig. 2c), validating stromal localization.

### Real-time monitoring of light-induced chloroplast stromal pH dynamics in leaf discs

Using a recently developed and validated custom LED illumination device compatible with CLSM (Elsässer et al., 2020; Zheng et al., 2025; Fig. 3a) to enable biosensor imaging in the presence and absence of photosynthetic activity, we recorded the fluorescence response of each stromal sensor in Arabidopsis leaf discs using a standardized dark–light–dark transition timelapse protocol (10 min per phase). We used three actinic light intensities (25, 200 and 850 µE; 1 µE = 1 µmol photons m^-2^ s^-1^; Fig. 3b). Ratios were normalized to baseline and are shown such that an increase indicates alkalinization. For Lime, the fluorescence at around 400 nm excitation was low and could not be reliably discriminated from the background. Hence the fluorescence intensity after 488 nm excitation is shown, relative to its own pre-illumination level, F/F_0_. At all three light intensities, illumination triggered a rapid rise in pH, consistent with light-driven stromal alkalinization, followed by a decline once the light was switched off. At 25 µE, pH rose quickly after light-on and approached a steady state. At 200 µE and, even more prominently, at 850 µE, all four sensors showed a transient alkaline overshoot, peaking shortly after the onset of illumination before settling into a signature similar to that at lower light intensities. Such an overshoot was previously reported for cpYFP (Elsässer et al., 2020; Zheng et al., 2025). While the overall signatures of the four sensors were similar, they differed in several aspects that were largely consistent with their different *in vitro* properties (Fig. 3b). Lime produced the largest response at all three light intensities (Fig. 3b; Fig. S3b), but since it was analyzed intensiometrically, the response amplitude strongly depends on the basal sensor expression level. cpYFP and pHluorin2 responded with similar amplitudes at 25 and 200 µE, and cpYFP stood out at 850 µE, where its overshoot was most pronounced. pHGFP showed the smallest changes, with a slow rise during illumination and little dependence on light intensity, although its AUC did not differ significantly from that of cpYFP and pHluorin2 at 25 and 200 µE (Fig. S3b). When the fluorescence intensity of the four sensors in the two excitation channels was compared under the same imaging settings (Fig. S3a), pHluorin2 was by far the brightest sensor in both excitation channels. Its signal was ∼8-fold higher than that of pHGFP at 405 nm and ∼2-fold higher than that of Lime at 488 nm.

**Figure 3.**
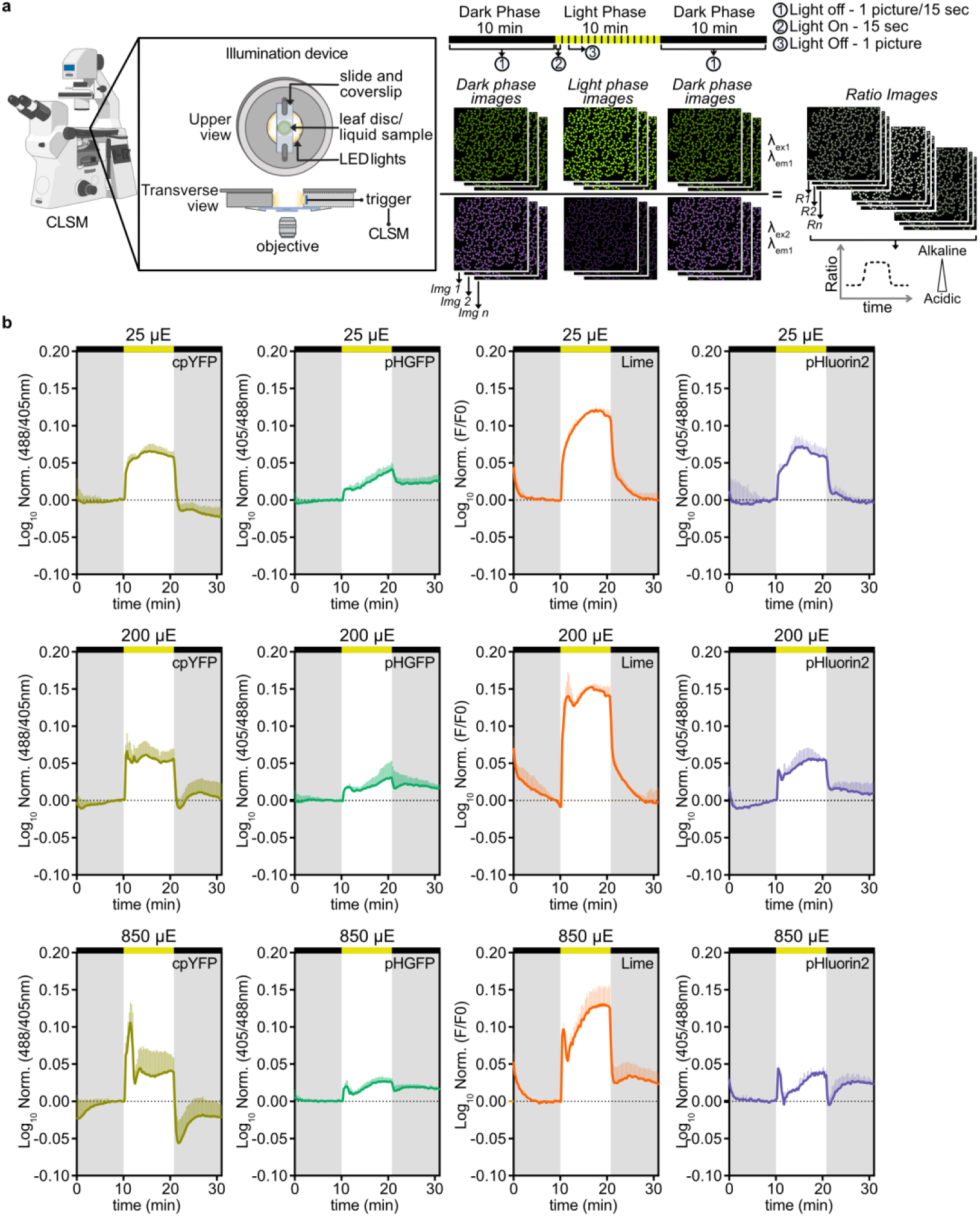
Light-induced responses of four stromal pH sensors in Arabidopsis leaf discs. **a.** Experimental set-up: leaf discs mounted in a custom LED illumination device on a CLSM, and the dark–light–dark protocol (10 min per phase). Dual-excitation images (λ_ex1_ = 488 nm, λ_ex2_ = 405 nm; emission 508–535 nm) were acquired every 15 s and converted to ratio images. Confocal Microscope and device cartoons were created in BioRender. Schwarzlander, M. (2026) https://BioRender.com/8ypyqho). **b.** Normalized ratios of cpYFP (488/405), pHGFP (405/488), pHluorin2 (405/488) and Lime (F/F_0_ of the 488 nm-excited fluorescence) at 25, 200 and 850 µE (rows; 1 µE = 1 µmol photons m^−2^ s^−1^). Each trace was normalized to the mean of the five time points before illumination and log_10_-transformed; an increase indicates alkalinization. The bar above each trace shows the illumination phase (black, dark; yellow, light). Lime is intensiometric and was therefore normalized to its own 488 nm fluorescence before illumination instead of to a ratio. Lines, mean; shaded bands, ± SD; n = 3 biological replicates (leaf discs from independent plants).

For cpYFP, fluorescence in the 405 nm excitation channel was low and close to background of leaf tissue (Fig. S3a), which warrants caution when interpreting its ratio. This problem is circumvented by using the intensiometric analysis for Lime, at the cost of internal normalization, however. pHGFP showed a small response amplitude, whereas pHluorin2 combined a consistent *in vivo* response with high brightness in both excitation channels (Fig. 2b; Fig. S3a) and a response at acidic pH *in vitro* (Fig. 1c; Table S1). Based on these empirical criteria we selected pHluorin2 for subsequent experiments.

### Expression of pHluorin2 in the chloroplast stroma of *Chlamydomonas reinhardtii* and light-induced responses

To expand our toolbox for photosynthesis-associated *in vivo* pH analysis also to algae, we generated *Chlamydomonas reinhardtii* lines in the CC125 and UVM4 backgrounds stably expressing pHluorin2 in the chloroplast stroma (CC125, Fig. 4a; UVM4, Fig. S4). Selected transformants were characterized by 2D-EEM scans of cell suspensions (CC125, Fig. 4b; UVM4, Fig. S4a). In addition to the endogenous fluorescence including the signal from the photosynthetic pigments in wild-type cultures, the transformants showed characteristic fluorescence above 500 nm. pHluorin2 fluorescence filled the chloroplast interior, as revealed by CLSM analysis (CC125, Fig. 4c; UVM4, Fig. S4b), showing a distribution distinct from chlorophyll fluorescence, i.e. consistent with stromal localization, and was also detected in the pyrenoid in CC125 (Fig. 4c). We did not calculate Pearson’s correlation coefficients for Chlamydomonas because, different to Arabidopsis and other plants, its thylakoids form appressed and non-appressed regions but do not form distinct grana stacks, preventing the systematic discrimination of thylakoids and stroma within the resolution achievable by live cell CLSM.

**Figure 4.**
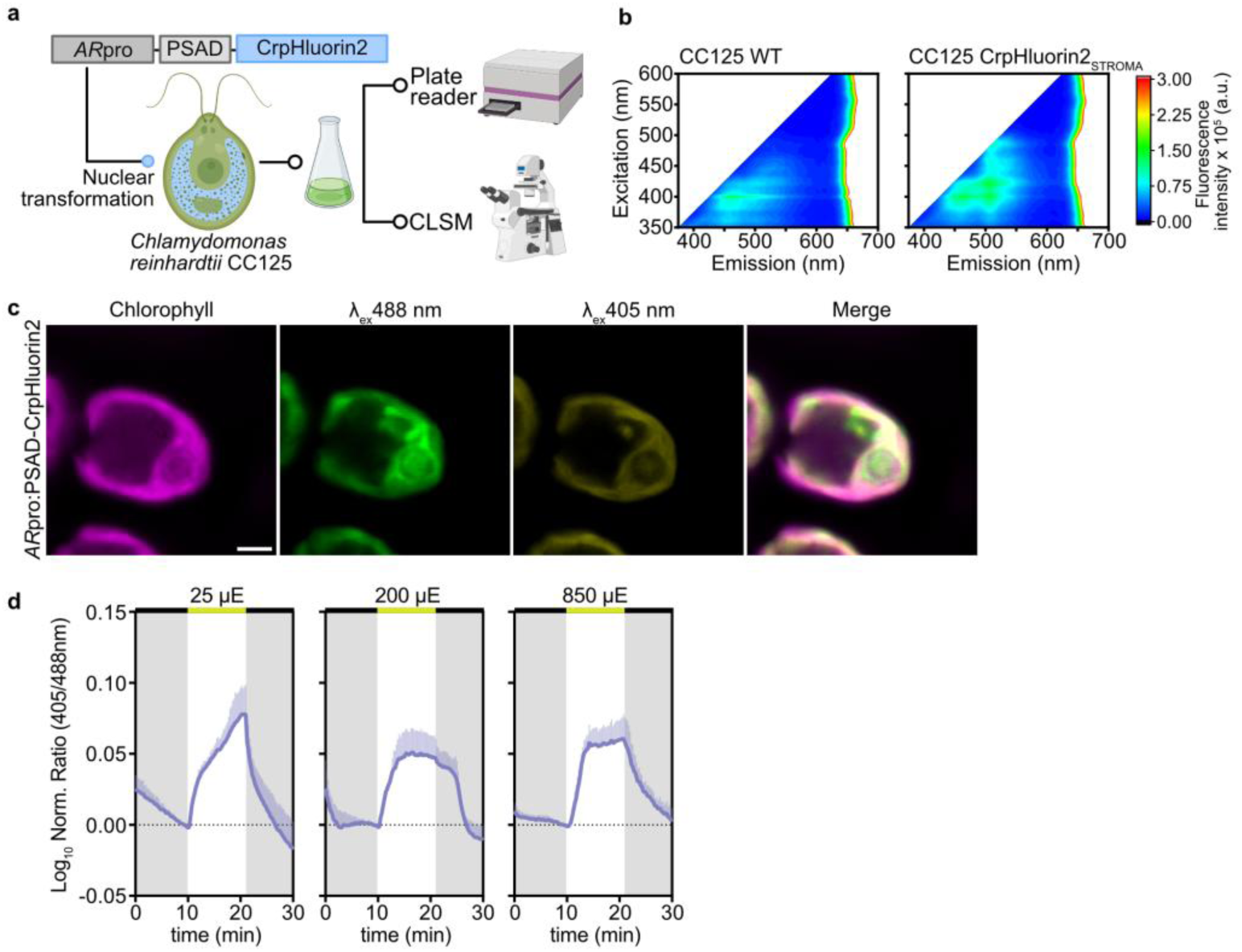
Stromal pHluorin2 in *Chlamydomonas reinhardtii*. **a.** Construct *AR*pro:PSAD-CrpHluorin2 used to express CrpHluorin2 (codon-optimized for Chlamydomonas) in the chloroplast stroma of *C. reinhardtii* CC125 by nuclear transformation and the analysis of the resulting cultures with a plate reader and a confocal microscope. *AR*pro, chimeric *HSP70A*–*RBCS2* promoter; PSAD, photosystem I subunit D (PsaD) chloroplast transit peptide. Plate reader, confocal microscope and Erlenmeyer flask icons were created with BioRender.com (Schwarzlander, M. (2026) https://BioRender.com/8ypyqho; https://BioRender.com/l3578yo). **b.** 2D-EEM of whole-culture suspensions of the wild type (WT) and the CrpHluorin2_STROMA_ line recorded on a plate reader; colour scale, fluorescence intensity (arbitrary units). **c.** CLSM images of a CC125 cell expressing CrpHluorin2_STROMA_: chlorophyll fluorescence (magenta), sensor fluorescence excited at 488 nm (green) and 405 nm (yellow), and the merge. Scale bar, 2 µm. **d.** Normalized 405/488 ratio (log_10_) of CrpHluorin2_STROMA_ in CC125 cells during a dark–light–dark protocol (10 min per phase) at 25, 200 and 850 µE. The bar above each trace shows the illumination phase (black, dark; yellow, light). Mean ± SD; n = 3 biological replicates (cultures grown in parallel from the same pre-culture). Cells were grown in TAP medium.

We then made use of the pH sensor to visualize photosynthesis-associated stromal pH dynamics by applying the same dark–light–dark protocol used for Arabidopsis (Fig. 4d). pH increased upon illumination and showed an overall response similar to Arabidopsis. That said, the pH signatures differed markedly in their characteristics from those observed in Arabidopsis. pH increased more gradually in response to the onset of illumination and did not show any alkalinization overshoot, even at 850 µE. At 25 µE, pH increased slowly throughout the light phase, whereas at 200 and 850 µE it rose faster and reached a *plateau*. After the light was switched off, pH declined at 25 and 850 µE. Remarkably, pH did not immediately decrease at onset of darkness after illumination at 200 µE but remained elevated for about two minutes to then decline. This behavior was consistently observed across experiments and in contrast to the signatures in Arabidopsis (Fig. 3b). The UVM4 background showed a similar light-dependent response at 200 and 850 µE, but no clear response at 25 µE (Fig. S4c). As such, the stromal pHluorin2 Chlamydomonas strains provide a tool to assess photosynthesis-associated pH dynamics in algae, and reveal both light-intensity dependence of the pH signatures, as well as distinct differences between the algal and the plant models during photosynthetic transitions.

### Light-induced pH changes reported by pHluorin2 in the thylakoid lumen

Monitoring pH dynamics in the lumen is of major interest, not only to understand ΔpH which requires an understanding of pH in both stroma and lumen, but also because of the regulatory role of lumen pH for NPQ and photosynthetic control. To that end, we generated stable Arabidopsis lines expressing pHluorin2 fused to the PsbQ1 transit peptide to facilitate luminal import (Fig. 5a). 2D-EEM confirmed expression (albeit at low level compared to stromal expression) and the presence of the characteristic fluorescence of the sensor *in vivo* (Fig. 5b). CLSM analysis showed pHluorin2 fluorescence exclusively in the chloroplasts, where it co-localized with chlorophyll fluorescence (Pearson’s r = 0.93; Fig. 5c), indicating localization to the thylakoid system. To discriminate presence of the biosensor in the lumen, from the possibility that the sensor is attached to the stromal-exposed face of the thylakoid membrane, we exploited the distinct pH response on either side of the thylakoid membrane to photosynthetic activity.

**Figure 5.**
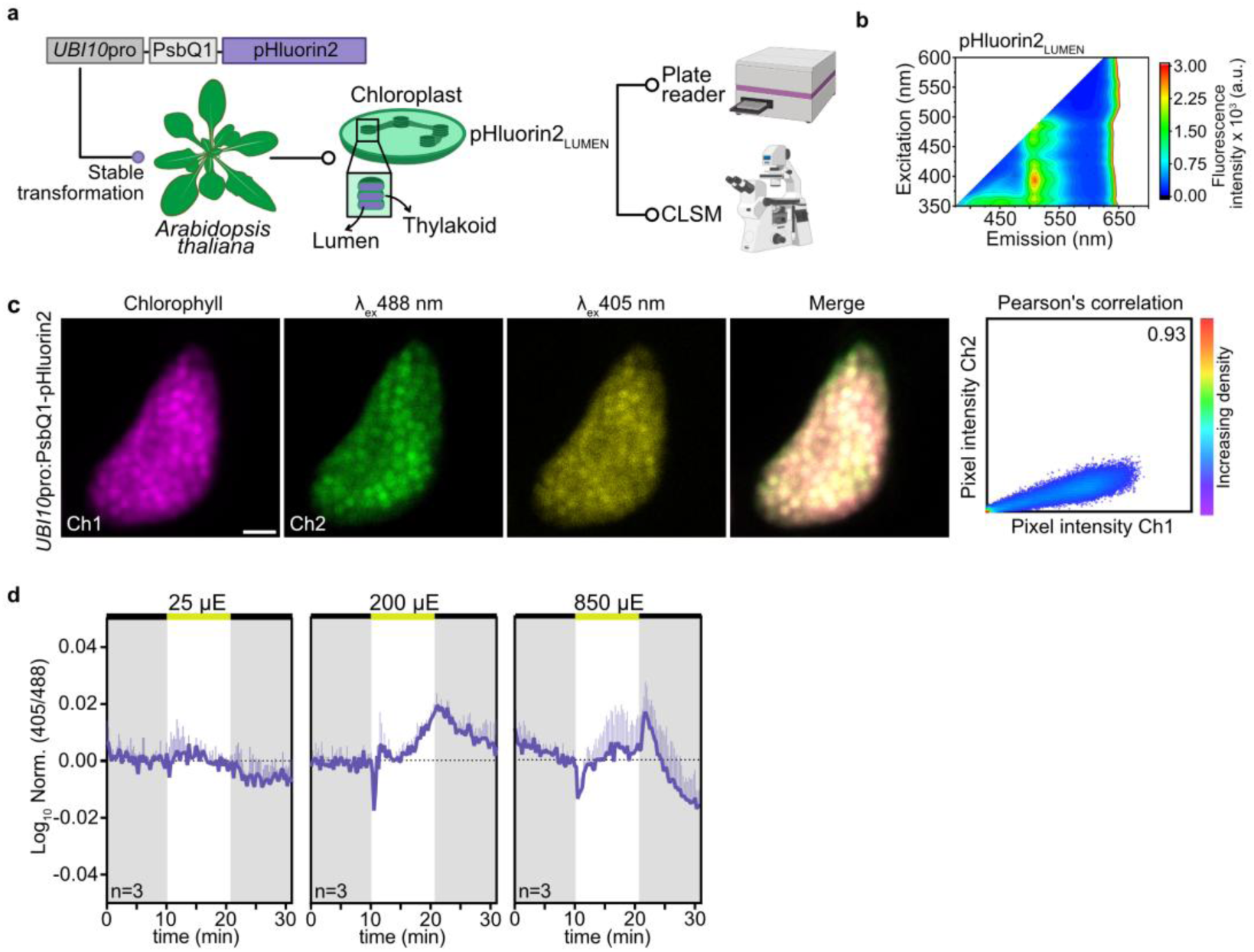
Thylakoid lumen-targeted pHluorin2 in Arabidopsis. **a.** Construct *UBI10*pro:PsbQ1-pHluorin2 used to generate a stable Arabidopsis line expressing pHluorin2 in the thylakoid lumen (pHluorin2_LUMEN_), and its analysis with a plate reader and a confocal microscope. PsbQ1, oxygen-evolving complex protein Q1 transit peptide. The plate reader and confocal microscope icons were created with BioRender.com (Schwarzlander, M. (2026) https://BioRender.com/8ypyqho). **b.** 2D-EEM of leaf discs of the pHluorin2_LUMEN_ line recorded on a plate reader; colour scale, fluorescence intensity (arbitrary units). **c.** CLSM of a mesophyll chloroplast showing chlorophyll fluorescence (Ch1, magenta), sensor fluorescence excited at 488 nm (Ch2, green) and at 405 nm (yellow), and the merge. Right, pixel-intensity scatter plot of Ch2 versus Ch1 (colour, pixel density); Pearson’s r = 0.93. Scale bar, 2 µm. **d.** Normalized 405/488 ratio (log_10_) of pHluorin2_LUMEN_ in leaf discs during a dark–light–dark protocol (10 min per phase) at 25, 200 and 850 µE. The bar above each trace shows the illumination phase (black, dark; yellow, light). Mean ± SD; n = 3 biological replicates (leaf discs from independent plants).

In leaf discs, illumination at 200 and 850 µE triggered a rapid, transient drop in pH, i.e. the opposite of what we observed for the stromal targeted biosensor. Indeed, this initial phase of the pH signature was qualitatively a mirror image of the stromal overshoot observed at the same intensity range (Fig. 3b), i.e. lumen acidified as the stroma alkalinized. Yet, unexpectedly the initial acidification event was followed by a recovery to pre-illumination pH and even above during the remainder of the light phase (Fig. 5d). At 25 µE, no clear response was resolved, which may reflect the small pH change expected at this intensity combined with the limited response range of pHluorin2 in the acidic range (Fig. 1c; Table S1). Although the amplitudes were small, the decrease at 200 and 850 µE occurred consistently across the replicates. As such the physiological data indicate that pHluorin2 is predominantly localized to the thylakoid lumen where it is able to respond to rapid proton accumulation at the onset of illumination. Yet, the subsequent sensor dynamics including the apparent loss of acidification are unexpected considering pH data from other approaches and will require further investigation before the luminal sensor data can be interpreted any further.

### Calibration of pHluorin2 for stromal pH estimation in isolated chloroplasts and live leaf tissue

To convert biosensor fluorescence changes into pH values, we calibrated stromal pHluorin2. Since our efforts to establish *in situ* pH calibration in intact leaves were unsuccessful, we opted for calibrations *in organello* where *in situ* pH can be externally controlled. Isolated Arabidopsis mesophyll chloroplasts were equilibrated in media of defined pH values in the presence of the ionophore nigericin to collapse transmembrane pH gradients (Fig. 6a). Excitation scans recorded by a multiwell fluorimeter showed the pH-dependent shift between the two excitation peaks of pHluorin2 (Fig. 6b) and yielded a sigmoidal relationship between fluorescence ratio and pH with an apparent pKa of 7.01. Analogous confocal imaging of the chloroplast samples delivered a consistent sigmoidal relationship with an apparent pKa of 7.10 (Fig. 6c). Both values were 0.2–0.3 pH units below the pKa of the purified recombinant protein (7.3; Table S1), highlighting the need for *in situ* calibration. The confocal titration was used to determine the relationship between 405/488 ratio and *in situ* pH (see Materials & Methods), and was applied to the subsequent estimates.

**Figure 6.**
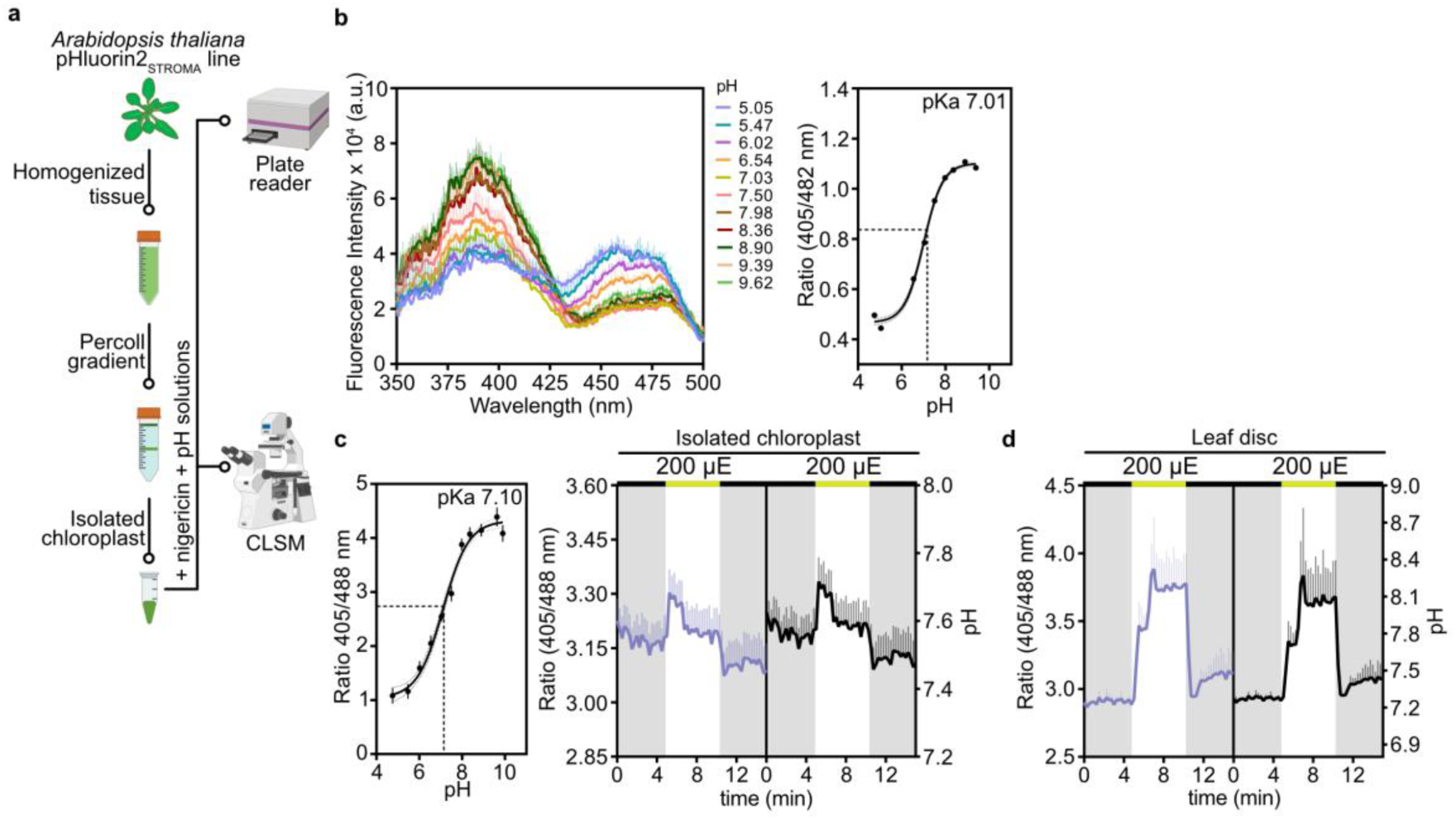
*In organello* calibration of stromal pHluorin2 and estimation of stromal pH. **a.** Workflow: chloroplasts were isolated from the stromal pHluorin2 line on a Percoll gradient, equilibrated at defined pH values with nigericin, and analyzed on a plate reader (b) and by confocal microscopy (c). Plate reader, confocal microscope, Falcon tube and eppendorf tube icons were created with BioRender.com (Schwarzlander, M. (2026) https://BioRender.com/8ypyqho, https://BioRender.com/l3578yo). **b.** Left, excitation spectra of pH-clamped chloroplasts (emission 510 nm; pH 5.05–9.62). Right, ratio of fluorescence excited at 405 and 482 nm versus pH, with a sigmoidal fit (apparent pKa 7.01). **c.** Left, 405/488 ratio of pH-clamped chloroplasts imaged by confocal microscopy versus pH, with a sigmoidal fit (apparent pKa 7.10); mean ± SD, five images per pH. Right, 405/488 ratio and pH of isolated chloroplasts during a dark–light–dark protocol (5 min per phase) under red light at 200 µE. The left axis shows the measured ratio (violet); the right axis shows the same trace converted to stromal pH (black) with the equation derived from the confocal calibration (non-linear scale). **d.** As in c (right) for leaf discs of the stromal pHluorin2 line: the left axis shows the 405/488 ratio (violet) and the right axis the corresponding stromal pH (black). Traces, mean ± SD; n = 3 chloroplast preparations or leaf discs. The bar above each trace shows the illumination phase (black, dark; yellow, light).

In isolated chloroplasts illuminated at 200 µE, pH increased upon illumination and decreased in the dark (Fig. 6c). This increase corresponded to a modest rise in stromal pH from ∼7.5 in the dark to ∼7.6 in the light, in the range of a previous report on isolated chloroplasts from spinach (Su & Lai, 2017). In leaf discs of the same biosensor line, the response was substantially larger: the stromal pH increased from ∼7.3 in the dark to ∼8.0 in the light and declined to ∼7.4 in the dark (Fig. 6d). These empirical values lie within the range of earlier estimates of light-induced stromal alkalinization obtained by independent methods (Heldt et al., 1973).

### Photosynthetic electron flow-associated generation of a proton motive force shapes stromal pH dynamics

To test whether the light-induced pH changes depended on photosynthetic activity, we used the photosystem II inhibitor 3-(3,4-dichlorophenyl)-1,1-dimethylurea (DCMU). DCMU blocks electron transfer from Q_A_ to plastoquinone, and thereby linear electron flow, and the associated proton translocation into the lumen (Fig. 7a). We applied DCMU to the leaf samples before starting the measurements using the standardized dark–light–dark regime (Fig. 7b). DCMU abolished the light-induced pH increase of stromal pHluorin2 in Arabidopsis leaf discs and diminished the drop in pH observed in luminal pHluorin2 in controls (Fig. 7b). In Chlamydomonas, DCMU also dampened the light-induced stromal pH increase, yet to a lesser extent than in Arabidopsis, indicating that the light-induced signal of pHluorin2 depended on photosynthetic electron transport and reflected photosynthesis-driven pH changes.

**Figure 7.**
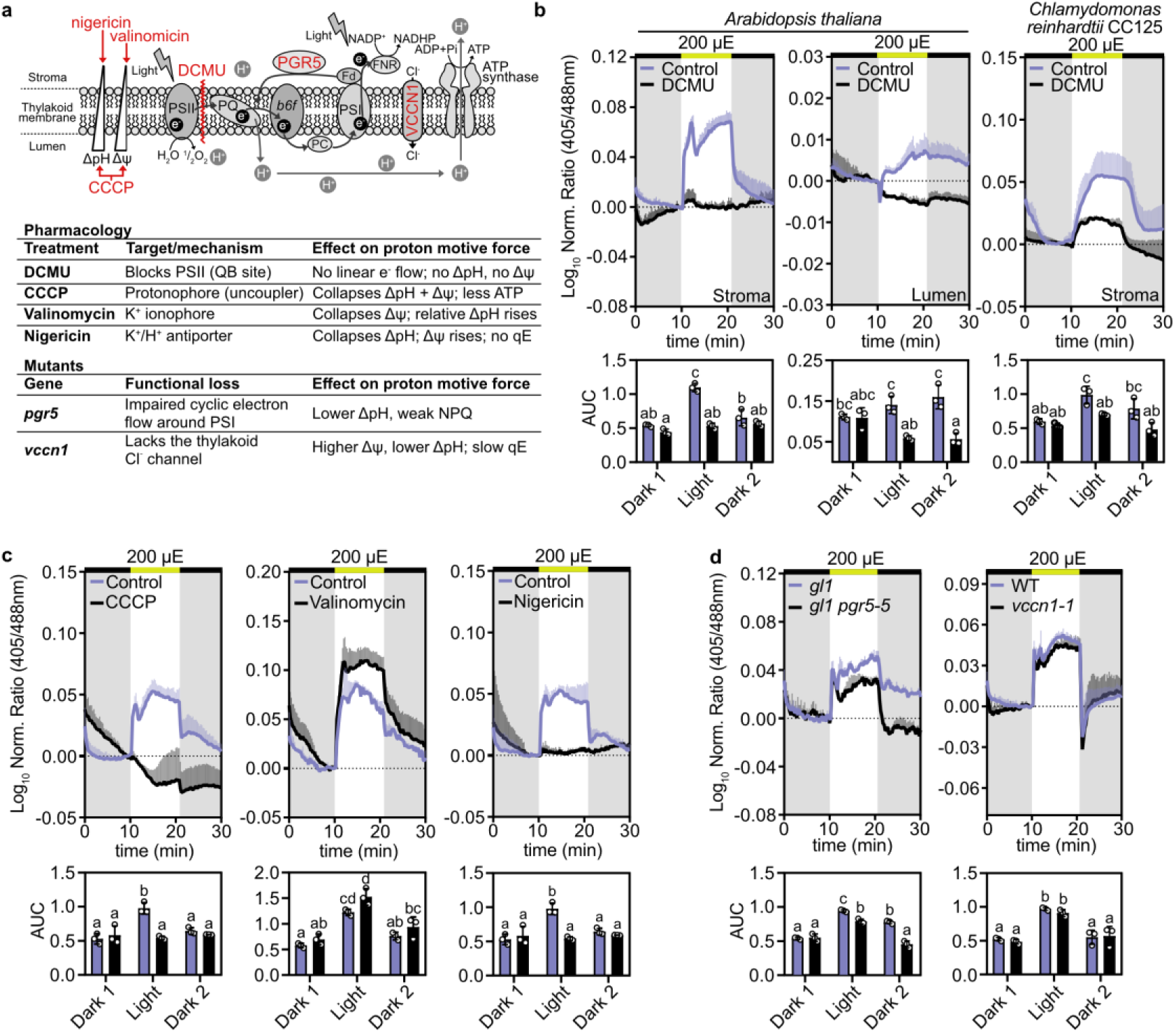
Dependence of the stromal pHluorin2 response on photosynthetic electron transport, the proton motive force, PGR5 and VCCN1. **a.** Scheme of the thylakoid membrane showing the sites of action of 3-(3,4-dichlorophenyl)-1,1-dimethylurea (DCMU, inhibiting photosystem II), nigericin (collapsing ΔpH), valinomycin (collapsing Δψ) and carbonyl cyanide *m*-chlorophenyl hydrazone (CCCP, dissipating ΔpH and Δψ), and the positions of PGR5 and VCCN1. The table summarizes the pharmacology of each treatment and the functional loss in each mutant, and the expected effect on the *pmf*. **b.** Normalized 405/488 ratio (log_10_) of pHluorin2 in untreated (violet) and DCMU-treated (black) Arabidopsis leaf discs expressing stromal (left) or luminal (middle) pHluorin2 and in *C. reinhardtii* CC125 cells expressing stromal pHluorin2 (right), during a dark–light–dark protocol (10 min per phase) at 200 µE. Leaf discs and cells were incubated with 20 µM DCMU for 30 min before the measurement; controls received the solvent (ethanol, 0.06% (v/v) final concentration). Below, area under the curve (AUC) of the normalized ratio in each phase (Dark 1, Light, Dark 2; baseline −0.05; −0.01 for luminal pHluorin2). **c.** As in b for Col-0 leaf discs expressing stromal pHluorin2, untreated (violet) or treated (black) with CCCP (10 µM), valinomycin (5 µM) or nigericin (2 µM) for 30 min; controls received the corresponding solvent (ethanol, 0.1% (v/v) final concentration, for CCCP; DMSO, 0.1% (v/v) final concentration, for valinomycin and nigericin). **d.** Stromal pHluorin2 in *gl1* (violet) and *gl1 pgr5-5* (black) (left) and in WT (violet, Col-0) and *vccn1-1* (black) (right) at 200 µE. In the AUC plots, bars show the mean ± SD and dots individual biological replicates. Different letters indicate significant differences among all groups within a panel (two-way ANOVA with Tukey’s multiple comparisons test, P < 0.05). The bar above each trace shows the illumination phase (black, dark; yellow, light). Traces, mean ± SD; n = 3 biological replicates (leaf discs from independent plants).

We next manipulated the *pmf* in leaf discs expressing stromal pHluorin2, using the protonophore carbonyl cyanide *m*-chlorophenyl hydrazone (CCCP), with the aim to dissipate both *pmf* components, i.e. ΔpH and Δψ (Heytler & Prichard, 1962). To isolate each component, we also used nigericin, an electroneutral K^+^/H^+^ exchanger that collapses ΔpH, and valinomycin, a K^+^ uniporter that collapses Δψ (Fig. 7a) (Pressman, 1976). Both CCCP and nigericin abolished the light-induced pH increase (Fig. 7c). By contrast, valinomycin did not abolish it, and the amplitude tended to be higher than in the control, although the difference in the area under the curve (AUC) of the normalized ratio was not significant (Fig. 7c). Thus, the stromal pH response depended on ΔpH, whether dissipated alone or together with Δψ, but not on Δψ alone.

### Lack of PROTON GRADIENT REGULATION 5 is associated with a dampened stromal pH response to light

The pharmacological treatments show that the stromal signal depends on photosynthetic electron flow and on ΔpH. However, inhibitors and ionophores act acutely on the whole membrane system and thus cannot pinpoint the mechanism of how the *pmf* is generated and how it partitions. We therefore exploited Arabidopsis mutants with established changes in photosynthesis-associated pH dynamics. PROTON GRADIENT REGULATION 5 (PGR5) is required for the PGR5-dependent pathway of cyclic electron flow around photosystem I, which returns electrons from ferredoxin to the plastoquinone pool and thereby increases proton translocation into the lumen and ΔpH formation without net NADPH production (Munekage et al., 2002, 2004; Suorsa et al., 2012). We equipped the *pgr5-5* background (Kobayashi et al., 2024) and its background line *gl1* (Col-0 with a mutation in the GLABROUS1 (GL1) gene) with the stromal pHluorin2 construct. At 200 µE, pH increased after light-on in both genotypes, followed by an alkalinization transient, as in Col-0 (Fig. 3). However, a smaller AUC in *pgr5-5* than in *gl1* indicated lower pH values during and after the light phase (Fig. 7d). Also at 25 µE, stromal alkalinization showed similar genotype-specific signatures, with *pgr5-5* showing a lower alkalinization than the control, albeit the difference in AUC did not reach statistical significance. At 850 µE the response signature was clearly distinct between the genotypes even though the AUC was unaltered (Fig. S5a). The lower stromal alkalinization in *pgr5-5* at 200 µE reflects a smaller light-driven proton uptake into the lumen due to the impairment of PGR5-dependent cyclic electron flow and showcases the usefulness of the pH sensing toolbox to assess the specific physiological consequences of genetic intervention with the photosynthetic machinery.

### Lack of the voltage-gated thylakoid chloride channel VCCN1 does not lead to detectable changes in the stromal pH response to light

As an additional mutant we chose *vccn1-1*, to manipulate *pmf* partitioning rather than *pmf* generation. VCCN1 is a voltage-dependent chloride channel of the thylakoid membrane that was proposed to conduct Cl^-^into the lumen at the onset of illumination; its loss shifts the *pmf* towards Δψ at the expense of ΔpH and slows the induction of non-photochemical quenching (Herdean et al., 2016; Fig. 7a). The pH response in a *vccn1-1* background (SALK_103612, Col-0 background; Herdean et al., 2016) equipped with stromal pHluorin2 was indistinguishable from that of the wild type control at both 200 µE (Fig. 7d) and 850 µE (Fig. S5b). Thus, loss of VCCN1 did not detectably change stromal pH dynamics under these conditions. The shift of the *pmf* towards Δψ in *vccn1-1* was reported to be moderate and the total *pmf* is higher than in the wild type (Herdean et al., 2016), so the resulting decrease in ΔpH may be too small to resolve with a stromal sensor that approaches saturation near pH 8 (Table S1), or it may be buffered in the stroma; the luminal sensor would be the more direct readout in this mutant.

Together, these experiments validate the utility of stromal pH sensing using pHluorin2 to monitor changes in photosynthesis associated pH dynamics.

## Discussion

### Comparison and establishment of pH sensors in plants

*In vivo* measurements of chloroplast pH remain scarce and have been constrained by the sensor or experimental system used. Dye-based measurements were restricted to isolated chloroplasts (Su & Lai, 2017), and the sensor PRpHluorin to non-photosynthetic plastids, because chlorophyll autofluorescence interferes with the measurement (Shen et al., 2013). cpYFP has since become a widely used tool to follow stromal pH in intact tissue and to resolve pH changes dynamically (Behera et al., 2018; Elsässer et al., 2020; Zheng et al., 2025; Uflewski et al., 2024), but its high pKa renders it insensitive in acidic compartments. A luminal pHluorin responded to light only in isolated chloroplasts (Yang et al., 2017). Tools that report pH in different subcellular compartments of intact cells are therefore still needed to resolve how the pH of stroma and lumen, and hence the *pmf*, is generated and regulated during photosynthesis.

To address this need, we compared four sensors *in vitro* and *in planta* (Fig. 1; Table S1). Their *in vitro* pKa values (8.4 for cpYFP, 7.3 for pHluorin2, 7.1 for pHGFP and 7.0 for Lime) determine which part of the physiological pH range each sensor resolves. cpYFP was the only sensor that retained a wide response range above pH 8 (5.1-fold between pH 7 and 8; 32% of its maximal response at pH 8), which makes it well suited to follow stromal alkalinization, whereas it and Lime responded too little between pH 5.5 and 7 for luminal measurements. In leaves, both cpYFP and Lime detected the light-induced alkalinization of the stroma with a clear response amplitude (Fig. 3). However, the weak signal of cpYFP after 405 nm excitation, which is close to the background of the leaf tissue, and the intensiometric readout of Lime mean that neither sensor provides a reliable internal normalization. A ratiometric readout makes the signal largely independent of expression level, tissue thickness and focal plane, which is what allows responses to be compared quantitatively between plants, genotypes and treatments. Without it, the amplitude of the cpYFP and Lime responses depends on the basal sensor level (Fig. 3; Fig. S3a), which limits their interpretation beyond a qualitative level. pHGFP, in turn, gave the smallest responses, with little dependence on light intensity. This could be explained by the small ratio change of pHGFP between pH 7 and 8 (1.5-fold, the lowest of the four sensors) and by its response being already ∼82% saturated at pH 8, so that stromal alkalinization falls on the flat upper part of its titration curve (Table S1). pHluorin2 combined the highest brightness with a ratiometric readout and remained responsive in the acidic range, and we therefore selected it for our subsequent *in vivo* studies.

### Physiological pH changes determined with pHluorin2

*In organello* calibration gave an apparent pKa of 7.0–7.1 for stromal pHluorin2 (Fig. 6), 0.2–0.3 units below that of the purified protein (7.3; Fig. 1c), showing that *in vitro* titrations cannot be transferred directly to the organelle. With this calibration, the stromal pH of isolated chloroplasts rose from ∼7.5 in the dark to ∼7.6 in the light, whereas in leaf discs it rose from ∼7.3 to ∼8.0. Published values depend on the technique, the species and the preparation. Classical measurements on isolated chloroplasts gave ∼7.0–7.2 in the dark and ∼7.8–8.0 in the light (Heldt et al., 1973; Werdan et al., 1975). BCECF in isolated spinach chloroplasts gave 7.32 in the dark and 7.55 in the light (Su & Lai, 2017), and in isolated Arabidopsis chloroplasts ∼7.4 in the dark and ∼7.7–7.8 in the light (Aranda Sicilia et al., 2021). Taken together, these studies span ∼7.0–7.4 in the dark and ∼7.55–8.0 in the light. The values obtained in this study are within or close to these ranges, which indicates that the *in organello* calibration of pHluorin2 yields physiologically plausible pH values. Two aspects limit the precision of the absolute values. First, a pH of ∼8.0 lies close to the upper end of the response range of pHluorin2 (78% of its maximal response at pH 8; Table S1), where the ratio changes less per pH unit, so that values for illuminated leaf discs are less precise than those in the dark. Second, the calibration was obtained in isolated chloroplasts and applied to intact tissue without an independent *in situ* calibration. We therefore regard the absolute values as estimates, whereas the direction and kinetics of the pH changes do not depend on the calibration. The larger response in tissue than in isolated chloroplasts may reflect metabolite exchange with the cytosol and mitochondria and the ATP demand of the intact cell, which are retained in leaf discs but altered upon chloroplast isolation. A key advantage of the approach presented here is its temporal resolution. Whereas classical estimates were obtained by rapid fractionation of chloroplasts after minutes of illumination, confocal imaging of intact tissue offers a time resolution of seconds and many measurements per experiment, and therefore resolves fast events such as the overshoot at moderate and high light (Fig. 3; Elsässer et al., 2020; Zheng et al., 2025).

### Lumen pH

In the thylakoid lumen, pHluorin2 reported a rapid, transient drop in pH within the first minute of illumination, which coincided with the stromal overshoot and was opposite in direction. This is consistent with fast proton loading of the lumen at the onset of illumination, before proton efflux through the ATP synthase and metabolic sinks have adjusted, and the drop was diminished by DCMU (Fig. 7b), indicating that it depends on photosynthetic electron transport. The subsequent recovery of the pH to, and above, the pre-illumination level during continued light was, however, unexpected considering the sustained lumen acidification inferred from other approaches (Rumberg & Siggel, 1969; Kramer et al., 1999). The opposite direction relative to the stromal sensor and the co-localization with chlorophyll (Pearson’s r = 0.93) indicate that the sensor is predominantly luminal. Its fluorescence is low, however, and we cannot exclude that part of the signal originates from sensor molecules at the stromal face of the thylakoid membrane. The luminal system therefore requires critical validation before it can be used as a tool, for example by a lumen-specific calibration in isolated thylakoids and by tests in mutants with altered lumen acidification.

### pH responses in algae

Stromal pHluorin2 was functional in *Chlamydomonas reinhardtii*, which establishes the sensor as a tool to follow photosynthesis-associated pH changes in a unicellular alga, where pH is difficult to access by other means. In contrast to Arabidopsis, stromal pH in Chlamydomonas rose gradually without overshoot and remained elevated after the light was switched off, which indicates that the build-up and decline of stromal pH are regulated differently in the two organisms. Several features of Chlamydomonas could contribute to the gradual rise without an overshoot: flavodiiron proteins, which are absent from angiosperms and mediate fast O_2_ photoreduction in the alga (Chaux et al., 2017); a different balance between linear and cyclic electron flow (Alric, 2010); and the pyrenoid-based carbon-concentrating mechanism (He et al., 2023). The ATP synthase could also contribute to the slow decline in the dark. In land plants, a redox switch in the γ-subunit inactivates the enzyme in the dark, whereas in Chlamydomonas the γ-subunit redox domain eludes this regulation (Lebok & Buchert, 2024), so that the enzyme may stay active in the dark and sustain a proton gradient through ATP hydrolysis (Rappaport et al., 1999; Nawrocki et al., 2015). These hypotheses can now be addressed with mutants of the respective components. As the cells were grown mixotrophically in TAP medium, the contribution of acetate metabolism and of the crosstalk between chloroplasts and mitochondria to these kinetics remains to be tested under photoautotrophic conditions.

### Contribution of ΔpH, Δψ and cyclic electron flow to the stromal pH response

The pharmacological and genetic experiments show that the stromal signal is specific to the *pmf* and that the sensor distinguishes its components. DCMU, which stops proton pumping by linear electron flow, and nigericin and CCCP, which collapse ΔpH, abolished the light-induced rise, whereas valinomycin, which collapses only Δψ, preserved it. Valinomycin also served as a control: it perturbs the membrane potential but did not abolish the signal, so the loss of response with nigericin reflects the loss of ΔpH rather than a general membrane artifact. The amplitude of the trace with valinomycin tended to be larger than in the control, although the AUC did not differ significantly. This tendency would be consistent with a shift of the pmf towards ΔpH when Δψ is dissipated, because the loss of Δψ relieves the electrical backpressure on proton pumping and reduces proton efflux through the ATP synthase, so that more protons accumulate in the lumen and a larger fraction of the pmf is stored as ΔpH (Braun et al., 2007; Davis et al., 2017).

In *pgr5-5*, which is impaired in PGR5-dependent cyclic electron flow (Munekage et al., 2002, 2004; Kobayashi et al., 2024), stromal alkalinization was smaller than in *gl1* at 200 µE, whereas the AUC did not differ significantly at 25 and 850 µE (Fig. 7d; Fig. S5a). This indicates that cyclic electron flow contributes to ΔpH formation, and hence to stromal alkalinization, at moderate light. The partial reduction is consistent with the lower *pmf* of *pgr5* mutants (Avenson et al., 2005), which results from the loss of PGR5-dependent cyclic electron flow rather than from proton leakage (Yamamoto & Shikanai, 2020) and reaches ∼70–75% of the wild-type level at high light (Degen et al., 2026). After the light was switched off, stromal pH remained higher in *gl1* than in *pgr5-5*. A smaller *pmf*, and therefore less ΔpH to dissipate, could explain the faster return in the mutant, but the origin of the persistent post-illumination pH in *gl1* and a possible influence of the genetic background remain to be tested. The similar AUC at 850 µE, despite the different response signature (Fig. S5a), may reflect a larger contribution of linear electron flow to proton uptake at high light, or limits imposed by the working range of the sensor. By contrast, *vccn1-1* did not differ from Col-0, which suggests that the partitioning of the *pmf* controlled by VCCN1 (Herdean et al., 2016) has no detectable effect on stromal pH under these conditions. This could reflect compensation by other ion fluxes or the limited sensitivity of the stromal readout.

In summary, we developed pHluorin2 for monitoring pH changes in living photosynthetic cells, with an unprecedented compartment specificity and time resolution, although the luminal readout still requires critical validation. In the stroma and thylakoid lumen of Arabidopsis and in the chloroplast of *Chlamydomonas*, pH dynamics during dark–light transitions are fast and compartment-specific. These dynamics differ between organisms and depend on ΔpH and electron flow. These observations raise new questions about how the *pmf* is built, partitioned and used, and our toolkit can now be used to address them.

## Material and methods

### Constructs

Binary and expression vectors were generated by modular Type IIS Golden Gate assembly. For stromal expression in Arabidopsis, the coding sequences of pHGFP, pHluorin2 and Lime were cloned into pL0_CDS2Stop, and the promoters of *ASPARTIC PROTEASE A1* (*APA*) and *UBIQUITIN 10* (*UBI10*) into pL0_NT1. The transit peptides of NADPH-dependent thioredoxin reductase C (NTRC; AT2G41680; stroma) and oxygen-evolving complex protein Q1 (PsbQ1; AT4G21280; thylakoid lumen) were cloned into pL0_SP. Multi-part assembly into the level 2 binary vector pL2 yielded the constructs *UBI10*pro:NTRC-pHGFP, *APA*pro:NTRC-pHluorin2, *APA*pro:NTRC-Lime and *UBI10*pro:PsbQ1-pHluorin2, each carrying the *HSP18.2* terminator and a hygromycin B resistance cassette. The stromal cpYFP line (*35S*pro:TKTP-cpYFP) was described previously (Behera et al., 2018). For recombinant protein expression in *Escherichia coli*, the coding sequences of cpYFP, pHGFP, pHluorin2 and Lime were transferred via pL0_CDS1Stop into pET28GG, which adds a C-terminal 6×His tag. For nuclear transformation with chloroplast targeting in *C. reinhardtii*, a codon-optimized pHluorin2 sequence (CrpHluorin2) was assembled with the *Chlamydomonas* MoClo system (Crozet et al., 2018; Niemeyer & Schroda, 2022) into the construct *AR*pro:PSAD-CrpHluorin2, which contains the chimeric *HSP70A–RBCS2* promoter (*AR*pro; also known as pAR in the MoClo toolkit), the photosystem I subunit D (PSAD) transit peptide, RBCS2 introns to enhance expression, a MultiStop sequence, the RPL23 terminator and an aadA spectinomycin resistance cassette.

### Plant material and growth conditions

All *Arabidopsis thaliana* lines were in the Columbia-0 (Col-0) accession, except for *pgr5-5*, which was generated in the *gl1* background (Col-0 carrying a mutation in *GLABROUS1*) and therefore compared with *gl1*. The *pgr5-5* allele is a CRISPR–Cas9 knockout with a 14-bp deletion in the first exon of *PGR5* (Kobayashi et al., 2024), and *vccn1-1* is a T-DNA insertion line in Col-0 (SALK_103612; Alonso et al., 2003; Herdean et al., 2016). The stromal pHGFP, pHluorin2 and Lime lines (*UBI10*pro:NTRC-pHGFP, *APA*pro:NTRC-pHluorin2 and *APA*pro:NTRC-Lime), the luminal pHluorin2 line (*UBI10*pro:PsbQ1-pHluorin2) and the stromal pHluorin2 lines in *gl1*, *pgr5-5* and *vccn1-1* were generated by *Agrobacterium*-mediated transformation (see below). Seedlings were germinated on plates for 10 days, transplanted into pots with Einheitserde Profisubstrat (Einheitserde Werkverband e.V., Germany), covered with transparent domes for 24 h and grown under a 16-h light/8-h dark photoperiod (22 °C/18 °C) at 60% relative humidity and 120 µE (1 µE = 1 µmol photons m^-2^ s^-1^) from white LEDs. Plants were watered with tap water every two to three days and used at 4–5 weeks of age. Homozygous transgenic lines were identified by segregation analysis of T3 progeny.

### Plant transformation and selection

Binary vectors were introduced into *Agrobacterium tumefaciens* strain AGL1 by electroporation (Mersereau et al., 1990), and *A. thaliana* plants were transformed by floral dip (Zhang et al., 2006). Harvested seeds were surface-sterilized with 80% (v/v) ethanol and sown on half-strength Murashige and Skoog (MS) medium with 0.1% (w/v) sucrose, 0.05% (w/v) MES (pH 5.8) and 0.8% (w/v) plant agar. Seeds were stratified for 48 h at 4 °C and plates were incubated at 18 °C under a 16-h light/8-h dark photoperiod at 120 µE. Primary (T1) transformants were selected on 40 µg mL^-1^ hygromycin B, and transgene expression was verified by CLSM in leaves of 2-week-old plants.

### Chlamydomonas reinhardtii growth and transformation

*C. reinhardtii* strains CC125 and UVM4 were grown in Tris-acetate-phosphate (TAP) medium at 25 °C under a 16-h light/8-h dark photoperiod at 60 µE. Both strains were transformed by electroporation with the same protocol, using the linearized plasmid carrying the construct *AR*pro:PSAD-CrpHluorin2 (see Constructs). Cells from 200 mL of an exponentially growing culture were harvested (3,500 rpm, 10 min, 16 °C) and resuspended in 80% TAP with 40 mM sucrose to 1 × 10^8^ cells mL^-1^. After incubation of 300 µL of cells with 1 µg of linearized plasmid for 20 min at 16 °C, 250 µL were electroporated in 1-mm cuvettes with a Gene Pulser II and Pulse Controller Plus (Bio-Rad) at 1,000 V and 10 µF (UVM4) or 25 µF (CC125). Cells were recovered for at least 20 min at 16 °C, mixed with TAP containing 40 mM sucrose and 20% (w/v) starch, and plated on TAP agar with spectinomycin (150 µg mL^-1^ for UVM4 and 300 µg mL^-1^ for CC125). Plates were kept in the dark overnight and then transferred to light. Transformants were first screened for pHluorin2 fluorescence with a fluorescence stereomicroscope. Positive transformants were confirmed by CLSM, and selected lines were characterized by whole-culture 2D-EEM scanning.

### Recombinant protein expression, purification and pH titration

*E. coli* BL21 (DE3) Rosetta cells were transformed with the pET28GG plasmids by heat shock (Froger & Hall, 2007) and grown in LB medium with kanamycin (50 µg mL^-1^) at 37 °C to an OD_600_ of 0.6. Expression was induced with 0.5 mM IPTG overnight at 20 °C. Cells were lysed in binding buffer (50 mM Tris-HCl pH 7.5, 100 mM NaCl, 0.5 mM MgCl_2_) with 1 mg mL^-1^ lysozyme, 0.1 mg mL^-1^ DNase I and EDTA-free protease inhibitor cocktail and sonicated. The cleared, filtered lysate was loaded onto a Ni-NTA gravity column, washed with 20 and 40 mM imidazole, and proteins were eluted with 250 mM imidazole, desalted into storage buffer (50 mM MES-KOH pH 7.0, 10% v/v glycerol) with Zeba Spin Desalting Columns (7K MWCO; Thermo Fisher Scientific) and stored at −80 °C. Protein concentrations were determined by Bradford assay against BSA standards. For pH titrations, each protein (0.1 mg mL^-1^) was diluted in a universal buffer (25 mM MES, 25 mM HEPES, 25 mM Tris, 25 mM CAPS and 100 mM NaCl) adjusted to 15 pH values from 3.99 to 10.98, and 200 µL were loaded in quadruplicate into a 96-well plate. The pH in each well was determined after the spectroscopic measurement and these empirical values were used for analysis. Spectra were recorded on a CLARIOstar microplate reader (BMG LABTECH): excitation spectra (350–500 nm) at 510 nm emission, and emission spectra from 475 to 600 nm (405 nm excitation) and 500 to 600 nm (482 nm excitation; Fig. S1). Blank buffer was subtracted. Points at the most acidic and alkaline pH, at which fluorescence was quenched (Martinière et al., 2013) or the proteins were denatured (Ward & Bokman, 1982), were excluded. The ratio of the excitation intensities at 482 and 400 nm (cpYFP, Lime) or 400 and 482 nm (pHGFP, pHluorin2) was plotted against pH and fitted with a sigmoidal function to obtain the apparent pKa, Hill coefficient and fold changes (Table S1).

### *In vivo* fluorescence spectral profiling

Excitation–emission matrices (2D-EEM) were recorded from leaf discs of 4- to 5-week-old plants submerged in 200 µL of assay buffer (10 mM MES-KOH pH 5.8, 10 mM MgCl_2_, 10 mM CaCl_2_, 5 mM KCl) in clear 96-well plates, from the top on a CLARIOstar plate reader. Excitation was stepped from 350 to 600 nm in 5-nm increments, and emission was recorded from 25 nm above the excitation wavelength to 700 nm. The gain was set automatically to 80% of the detector range on one well and kept fixed for all genotypes. Three leaf discs per genotype were measured and a representative scan is shown. Col-0 leaf discs were measured in parallel as autofluorescence reference, and no background was subtracted. Chlamydomonas cultures (200 µL) were scanned with the same settings (three replicates per strain).

### Confocal microscopy

Subcellular localization was determined on a Zeiss LSM 980 equipped with an Airyscan 2 detector (Super-Resolution mode) using a Plan-Apochromat 63×/1.40 Oil DIC M27 objective and Immersol 518F immersion oil. Leaf discs from 4- to 5-week-old plants were mounted in perfluorodecalin and imaged in the abaxial mesophyll or epidermal cell layer. Biosensors were excited sequentially at 405 and 488 nm and emission was collected between 495 and 550 nm; chlorophyll fluorescence was excited at 405 nm and collected at 655–735 nm. Laser power and detector gain were set to the minimum needed to avoid saturation and photobleaching, and Airyscan images were processed in ZEN (Carl Zeiss). Images were acquired at 512 × 512 pixels with a pixel size of 26.3 nm. Colocalization between the 488 nm-excited sensor channel and the chlorophyll channel was quantified in Fiji with the Coloc 2 plugin as the Pearson’s correlation coefficient within a region of interest covering the whole chloroplast; pixel-intensity scatter plots were generated in Origin (OriginLab).

### Biosensor imaging and light–dark regimes

Leaf discs from 4- to 5-week-old plants were mounted between two glass coverslips in 10 µL of perfluorodecalin. For imaging, Chlamydomonas cells were pre-cultured for 3 days, transferred to fresh TAP medium and used 48 h (CC125) or 24 h (UVM4) later, and CC125 cells were immobilized on poly-lysine-coated coverslips. Plants and algae were dark-adapted for at least 1 h before sample preparation. Imaging was performed on a Zeiss LSM 980 confocal microscope with a Plan-Apochromat 10×/0.45 M27 objective for the adaxial mesophyll layer of leaf discs and a 25× objective for Chlamydomonas cells. Actinic light was delivered by the stage-integrated illumination system described by Zheng et al. (2025). Biosensors were excited sequentially at 405 and 488 nm and emission was collected from 508 to 535 nm with a GaAsP-PMT detector (pinhole 120 µm). Autofluorescence and chlorophyll fluorescence were recorded under 405 nm excitation at 430–450 nm and 651–701 nm, respectively. Light–dark regimes consisted of three consecutive phases (dark, light, dark) of 10 min each under white LED light, or 5 min each under red LED light for the comparison with isolated chloroplasts. Light intensities are indicated in the figures.

### Pharmacological treatments

CCCP (10 µM), valinomycin (5 µM) and nigericin (2 µM) were applied to leaf discs by vacuum infiltration with a syringe in water containing the inhibitor, followed by a 30-min incubation before imaging. DCMU (20 µM) was applied without infiltration by placing leaf discs in a DCMU solution for 30 min; Chlamydomonas cells were incubated with 20 µM DCMU for 30 min in their TAP growth medium. CCCP was dissolved in ethanol, DCMU in 60% (v/v) ethanol, and valinomycin and nigericin in DMSO; controls received the corresponding solvent at the same final concentration (ethanol, 0.1% (v/v) for CCCP and 0.06% (v/v) for DCMU; DMSO, 0.1% (v/v)).

### Chloroplast isolation and *in organello* pH calibration

Intact chloroplasts were isolated from 3-week-old plants of the stromal pHluorin2 line and Col-0, grown as a dense lawn and dark-adapted overnight. All steps were carried out in pre-cooled isolation buffer (0.3 M sorbitol, 20 mM HEPES-KOH pH 8.0, 10 mM NaHCO_3_, 5 mM MgCl_2_, 5 mM EDTA). Tissue was homogenized with an Ultra-Turrax (IKA), filtered through gauze and centrifuged (1,470 × g, 5 min, 4 °C). The crude pellet was loaded onto a step gradient of 82% and 30% Percoll (with 0.3 M sorbitol, 20 mM HEPES-KOH pH 8.0, 5 mM MgCl_2_, 5 mM EDTA) and centrifuged at 1,940 × *g* for 10 min. Intact chloroplasts from the 30%/82% interface were washed in wash buffer (0.3 M sorbitol, 20 mM HEPES-KOH pH 8.0, 5 mM MgCl_2_), and their intactness was verified by CLSM. Chlorophyll content was determined in 80% (v/v) acetone with Arnon’s equation (Arnon, 1949), as described by Yan et al. (2020).

For calibration, equal amounts of chloroplasts (by chlorophyll) were pelleted and resuspended in 1 mL calibration buffer at the corresponding pH (50 mM KCl, 1 mM MgCl_2_, 1 mM CaCl_2_, 50 mM buffering compound and 2 µM nigericin; MES for pH 4.0–6.5, HEPES for pH 7.0–8.0 and CHES for pH 8.5–10.5) and incubated for ∼1 h at room temperature. For plate reader measurements, 200 µL were loaded in quadruplicate (pHluorin2 chloroplasts) or triplicate (Col-0 chloroplasts), excitation spectra were recorded at 510 nm emission, and the 405 nm and 482 nm excitation channels were measured (Fig. 6b). The pH of each well was re-measured and used for analysis, and the signal of Col-0 chloroplasts at the same pH was subtracted. For confocal calibration, an aliquot of each suspension was mounted in an 8-well chambered coverslip (#1.5 high-performance cover glass, 0.17 ± 0.005 mm, standard base; Cellvis, Mountain View, CA, USA) and imaged with a 25× objective (five images per pH). Intensities at 405 and 488 nm excitation were quantified with the Redox Ratio Analysis (RRA) software (Fricker, 2016), Col-0 background was subtracted, and the 405/488 ratio was fitted with a sigmoidal function (apparent pKa 7.10). Points at which sensor fluorescence was lost were excluded. The fitted curve was inverted to convert ratios to pH: pH = 7.105 − (1/0.7259) × log_10_ [(4.32 − 1.039)/(R – 1.039) − 1], where R is the 405/488 ratio.

To follow light-induced pH changes, chloroplasts were incubated in calibration buffer at pH 7.5 without nigericin, pelleted, and 10 µL of the pellet were mounted between two coverslips. Samples were imaged with a 25× objective during a 5-5-5 min dark–light–dark regime under red light (200 µE), with the acquisition and illumination settings described above; the same regime was applied to leaf discs of the stromal pHluorin2 line. Ratio traces were converted to pH with the calibration equation.

### Ratiometric analysis and statistics

Time series acquired during light–dark regimes were quantified with the RRA software (Fricker, 2016). Mean fluorescence intensities after 405 and 488 nm excitation were extracted for each frame, and a whole-frame ratio was calculated as 405/488 for pHGFP and pHluorin2 and 488/405 for cpYFP. Lime is intensiometric, so its 488 nm-excited intensity was used instead of a ratio. Each value was normalized to the mean of the five time points preceding light onset and log_10_-transformed. The area under the curve (AUC) was calculated with a baseline of −0.05 in all figures, over the whole time course in Fig. S3b and for each 10-min phase in Fig. 7, except for the luminal pHluorin2 traces, for which a baseline of −0.01 was used. Differences among sensors were tested by one-way ANOVA with Tukey’s multiple comparisons test; for Fig. 7, AUC values were compared by two-way ANOVA with Tukey’s multiple comparisons test. Biological replicates were leaf discs from different plants for Arabidopsis and three cultures grown in parallel from the same pre-culture for Chlamydomonas. Data are shown as mean ± SD. Curve fitting, statistics and plotting were performed in GraphPad Prism 8.0.1. Sample sizes are given in the figure legends.

## Author contributions

Conceptualization: APC, MH, KBB, MSchw

Methodology: APC, MH, PB, AE, KZ, LF, ME, SK, EW, MSchw

Investigation: APC, MH

Visualization: APC

Funding acquisition: MSchw, KBB

Project administration: MSchw, KBB

Supervision: MSchw, KBB, MHip, FB, MSchr, H-HK, UA

Writing – original draft: APC

Writing – review & editing: APC, PB, MH, MHip, FB, H-HK, UA, KBB, MSchw

## Acknowledgements

We thank Jens Appel (University of Kassel, Germany) for providing the pHluorin2 sequence and Johanna von Kielmansegg for assistance with cloning the plasmids for recombinant protein expression.

