## Supplemental Figures 1-5 for "Imaging photosynthetic pH dynamics in living plants and algae"

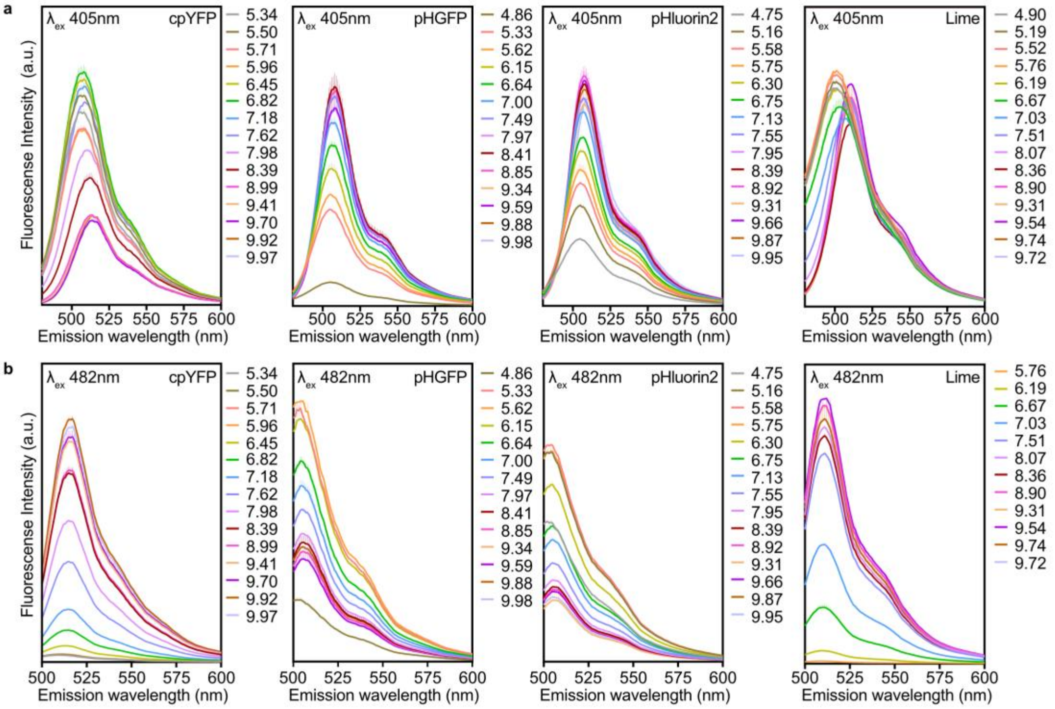


**Supplementary Figure 1 | Emission spectra of the purified sensors.**
**a, b.** Emission spectra of recombinant cpYFP, pHGFP, pHluorin2 and Lime at the indicated pH (colour code) with excitation at 405 nm (a) and 482 nm (b). Mean of n = 4 technical replicates (four wells of the same protein preparation). Fluorescence intensity is in arbitrary units.

| | Ratio | pKa | Response range | $n_H$ | Fold change, pH7-8 | Fold change, pH5.5-7 | % of max response at pH7 / 8 |
|---|---|---|---|---|---|---|---|
| **cpYFP** | 482/400 | 8.38 | 7.3 - 9.5 | 0.85 | 5.10 | n.d. | 6 / 32 |
| **pHGFP** | 400/482 | 7.12 | 5.8 - 8.4 | 0.74 | 1.54 | 2.3 | 45 / 82 |
| **Lime** | 482/400 | 7.03 | 6.4 - 7.7 | 1.38 | 2.04 | n.d. | 47 / 96 |
| **pHluorin2** | 400/482 | 7.30 | 5.9 - 8.5 | 0.78 | 1.67 | 2.1 | 37 / 78 |

**Supplementary Table 1 | *In vitro* characterization of the four pH sensors.**
Ratio orientation, apparent pKa, pH response range (interval between 10% and 90% of the maximal ratio response), Hill coefficient ($n_H$), fold change of the ratio between pH 7 and 8 and between pH 5.5 and 7, and percentage of the maximal response at pH 7 and 8, derived from sigmoidal fits to the titrations in Fig. 1c. n.d., not determined because the minimum ratio was approximately zero.

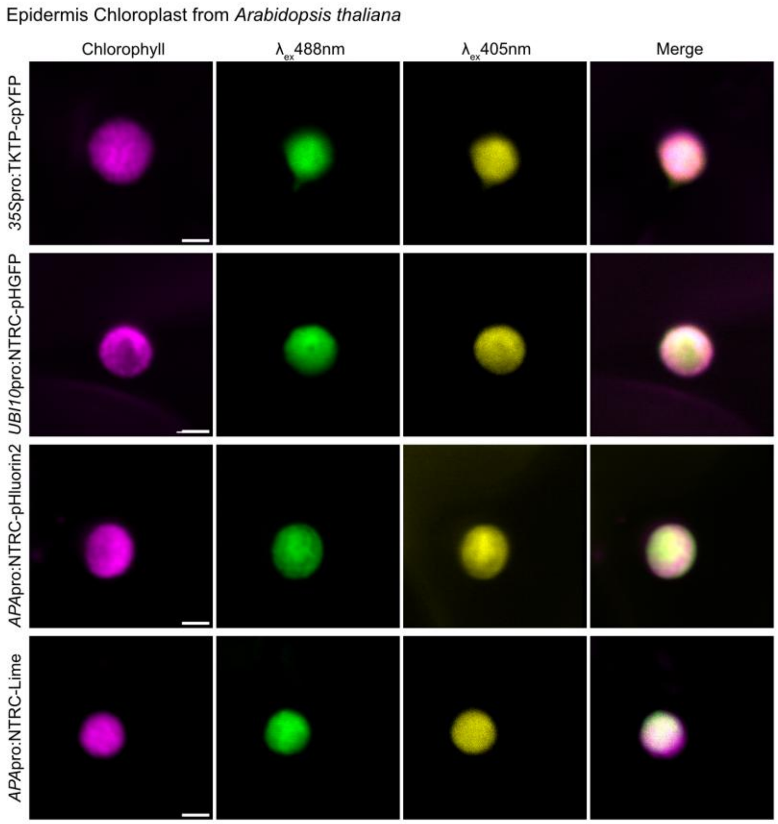


**Supplementary Figure 2 | Localization of the stromal sensors in epidermal chloroplasts.**
Confocal images of epidermal chloroplasts of Arabidopsis lines *35S*pro:TKTP-cpYFP, *UBI10*pro:NTRC-pHGFP, *APA*pro:NTRC-pHluorin2 and *APA*pro:NTRC-Lime: chlorophyll fluorescence (magenta), sensor fluorescence excited at 488 nm (green) and 405 nm (yellow), and the merge. Scale bars, 2 µm.

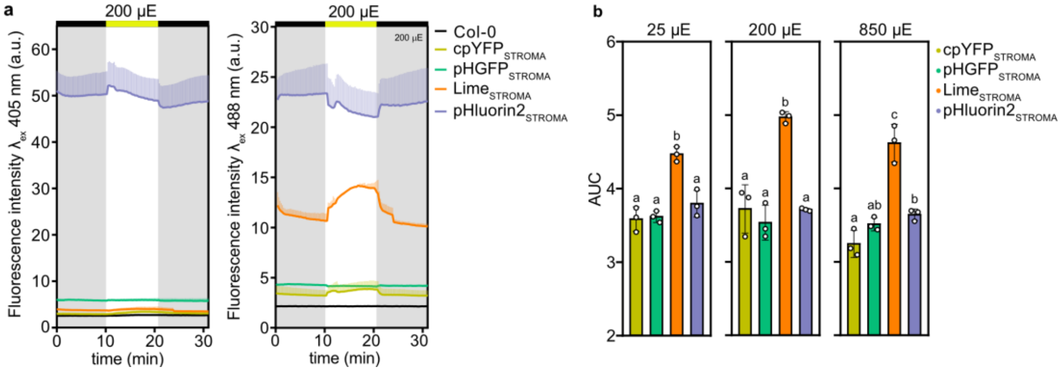


**Supplementary Figure 3 | Fluorescence intensity in the two imaging channels and AUC of the sensor responses.**

**a.** Fluorescence intensity (arbitrary units) after excitation at 405 nm (left) and 488 nm (right) in leaf discs of Col-0 and of the stromal cpYFP, pHGFP, Lime and pHluorin2 lines during a dark–light–dark protocol at 200 µE (bar as in Fig. 3). Lines, mean; shaded bands, ± SD. **b.** AUC of the traces in Fig. 3b, calculated over the whole time course with a baseline of −0.05, at 25, 200 and 850 µE. Bars, mean; error bars, ± SD; dots, individual biological replicates. Different letters indicate significant differences among all groups within a panel (one-way ANOVA with Tukey's multiple comparisons test, $P < 0.05$). n = 3 biological replicates (leaf discs from independent plants).

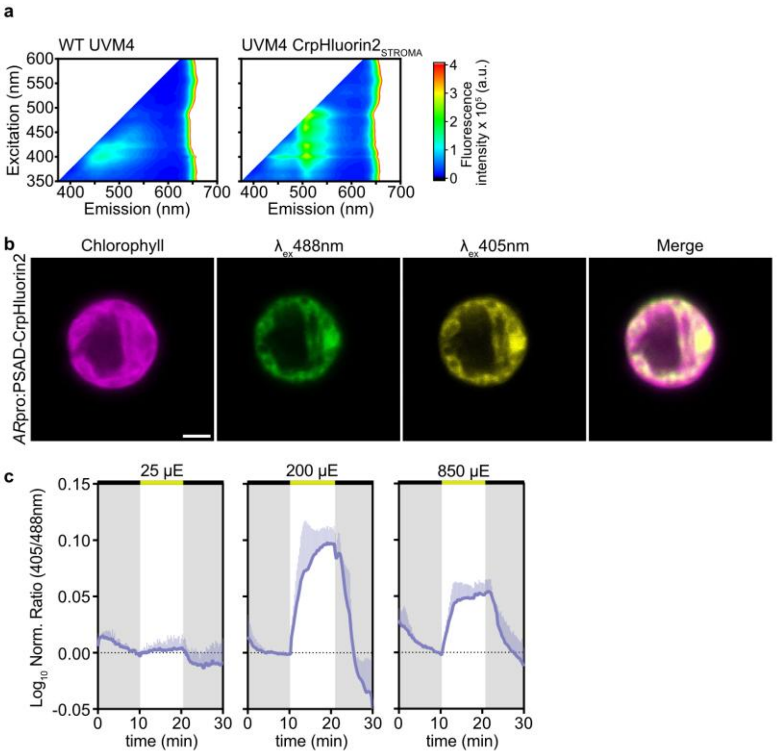


**Supplementary Figure 4 | Stromal pHluorin2 in *Chlamydomonas reinhardtii* UVM4.**
**a.** 2D-EEM of whole-culture suspensions of the UVM4 wild type (WT) and the CrpHluorin2$_{STROMA}$ line (excitation, y axis; emission, x axis; nm). **b.** Confocal images of a UVM4 cell expressing CrpHluorin2$_{STROMA}$: chlorophyll fluorescence (magenta), sensor fluorescence excited at 488 nm (green) and 405 nm (yellow), and the merge. Scale bar, 2 µm. **c.** Normalized 405/488 ratio ($\log_{10}$) during a dark–light–dark protocol (10 min per phase) at 25, 200 and 850 µE. The bar above each trace shows the illumination phase (black, dark; yellow, light). Mean ± SD; n = 3 biological replicates (cultures grown in parallel from the same pre-culture). Cells were grown in TAP medium.

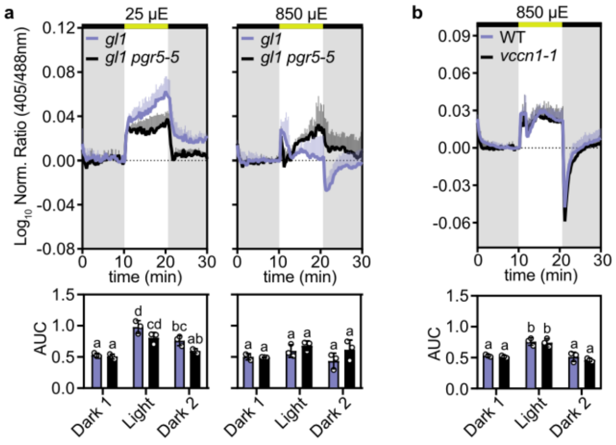


**Supplementary Figure 5 | Responses of stromal pHluorin2 in *pgr5-5* and *vccn1-1* at other light intensities.**
**a.** Normalized 405/488 ratio ($\log_{10}$) of stromal pHluorin2 in *gl1* (violet) and *gl1 pgr5-5* (black) leaf discs at 25 (left) and 850 µE (right), with the AUC of each phase below. **b.** As in a for WT (violet, Col-0) and *vccn1-1* (black) at 850 µE. Statistics, bars above traces and n as in Fig. 7.
